# A microprotein encoded by *FERMT3* modulates endothelial cell protein catabolism and induces p53-mediated cell cycle arrest and senescence

**DOI:** 10.64898/2025.12.03.692074

**Authors:** Manav Raheja, Beyza Güven, Witold Szymanski, Stefan Günther, Carsten Kuenne, Vladislav Rakultsev, Marta Segarra, Johannes Graumann, Ingrid Fleming, Mauro Siragusa

## Abstract

**Background:** Human endothelial cells express numerous microproteins (miPs) encoded by small open reading frames (smORFs) distributed throughout the genome, yet the biological functions of most remain unknown. This study set out to characterize a novel 69 amino acid miP encoded by a smORF located within the coding sequence of the FERM domain containing kindlin-3 transcript (miP-FERMT3).

**Methods:** Confocal microscopy was used to determine the subcellular localization of miP-FERMT3 in endothelial cells and its interaction partners were determined by mass spectrometry and immunoblotting. RNA sequencing identified transcriptional alterations induced by miP-FERMT3 overexpression. Cell proliferation and cell cycle stages were assessed by live cell imaging, EdU incorporation and flow cytometry, while senescence was examined by senescence-associated β-galactosidase staining, live cell imaging and RT-qPCR-based measurement of telomere length.

**Results:** In endothelial cells miP-FERMT3 localized mainly to centriole subdistal appendages, where it interacted with proteins involved in ubiquitin- and proteasome-dependent protein catabolism, including PSMD9, CUL2 and TRIM8. Consistent with these interactions, cells expressing miP-FERMT3 exhibited increased global protein ubiquitination, enhanced centrosomal neddylation and elevated proteasomal activity. MiP-FERMT3 also promoted the nuclear accumulation of p53, which subsequently repressed *FOXM1* expression, leading to the downregulation of genes required for cell-cycle progression and upregulation of genes involved in cell cycle inhibition, resulting in cell-cycle arrest. Cells expressing the miP also demonstrated multiple hallmarks of cellular senescence, including enlarged size, DNA damage, increased senescence-associated β-galactosidase activity, telomere shortening and paracrine pro-inflammatory activation of naïve endothelial cells. Analyses of independent murine and human transcriptomic and proteomic aging datasets further revealed that *FERMT3* expression and protein abundance increase with age.

**Conclusions:** These findings identify miP-FERMT3 as a novel regulator of protein catabolism and p53-dependent cell cycle arrest and cellular senescence in endothelial cells. Given the aging-associated upregulation of FERMT3 in mouse and human endothelial cells, increased miP-FERMT3 expression may contribute to the onset of vascular senescence as a hallmark of aging.

## INTRODUCTION

The genome contains thousands of open reading frames (ORFs) i.e., protein-coding sequences between in-frame start and stop codons. Breakthroughs in genomic and translational research have identified a surprisingly large number of small open reading frames (smORFs) within different classes of transcripts, including both coding and non-coding regions of annotated transcripts [1–4]. Although historically overlooked and thought to lack biological relevance, a substantial proportion of these elements are now recognized as functionally important. Some well-characterized examples reside in the 5′ untranslated regions (UTRs) of mRNAs, where they influence the translation efficiency of the downstream main coding sequence [5]. Beyond these regulatory roles, many smORFs encode short peptides of 100 or fewer amino acids, commonly referred to as microproteins (miPs) and significant efforts have improved their detection by mass spectrometry-based proteomic approaches [1, 2, 6]. Despite their relatively poor conservation across species [3, 5, 7, 8], these miPs have been shown to contribute to a range of fundamental processes, including DNA repair [1], calcium signalling [9–11], stress responses [12] and cell cycle control [13]. While some smORFs are ubiquitously expressed, others demonstrate high cell and tissue specificity [7].

Endothelial cells also express a large repertoire of previously unannotated smORF-encoded miPs [6, 7]. The endothelial smORFeome and microproteome vary across organs and inflammatory status [6]. Despite the limited sequence conservation between human and murine smORFs, we identified several smORFs whose expression was altered in IL-1β-treated human endothelial cells and in endothelial cells from a mouse model of endothelial dysfunction and accelerated atherogenesis. One of these smORFs is located within the coding sequence of the *FERM Domain Containing Kindlin 3* (FERMT3) transcript (ENST00000345728.10) but is translated in an alternative reading frame than the canonical ORF. This smORF is upregulated under inflammatory conditions *in vitro* and *in vivo* and encodes a 69 amino acid miP, hereafter referred to as miP-FERMT3. As this represents a previously uncharacterized miP, we investigated its function in human endothelial cells.

## RESULTS

### FLAG-miP-FERMT3 localizes to cytoplasm and centriole subdistal appendage

The FERMT3 smORF spans exons 5-7 of the human *FERMT3* gene (+chr 11:64,211,356-64,219,269) and is 210 bp long. (**Figure 1A**). The human miP-FERMT3 shares 69.6% homology with the mouse sequence and computational structure prediction (AlphaFold3) indicated that the miP adopts a predominantly linear, intrinsically disordered conformation (**Figure 1B**). To investigate the subcellular localization of miP-FERMT3, adenoviruses were generated to express a FLAG-tagged miP-FERMT3 fusion protein in endothelial cells. Adenoviral transduction induced an approximately 8-fold increase in *smORF-FERMT3* expression relative to control, which was comparable to levels observed in IL-1β-treated endothelial cell (**Figure 1C-D**). Next, FLAG immunofluorescence was combined with markers of specific subcellular organelles including the endoplasmic reticulum, Golgi apparatus, mitochondria, endosomes, lysosomes and centrosomes (**Figure 1E** and **Figure S1**). This approach revealed that FLAG-miP-FERMT3 was localized in the cytoplasm and accumulated near the Golgi apparatus where it colocalized with the centrosomal markers α-tubulin and γ-tubulin, as well as the centriole subdistal appendage markers ninein and CEP170.

**Figure 1.**
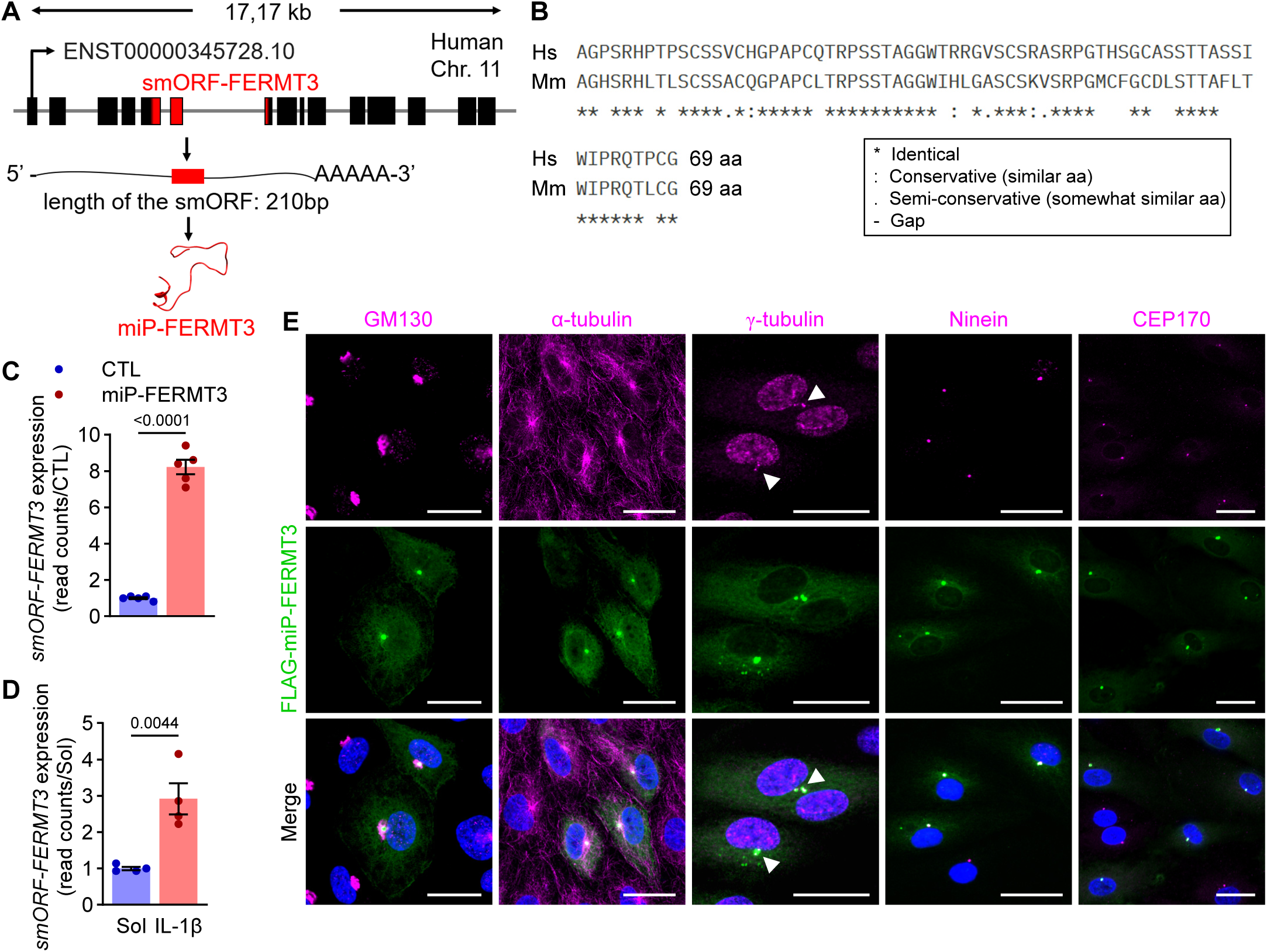
Location of smORF-FERMT3 and subcellular localization of miP-FERMT3. **(A)** Location of smORF-FERMT3 (red) in the *FERMT3* gene transcript and secondary structure prediction of miP-FERMT3 generated with AlphaFold3. **(B)** Sequence alignment of human and murine miP-FERMT3, generated with Clustal Omega. **(C)** *smORF-FERMT3* RNA expression (read counts) in endothelial cells expressing FLAG-miP-FERMT3 or miRFP647nano (CTL); n=5 independent cell batches (unpaired, two-tailed Student’s t-test). **(D)** *smORF-FERMT3* RNA expression (RiboTag read counts) in endothelial cells following treatment with solvent (Sol) or interleukin-1β (IL-1β); n=4 independent cell batches (unpaired, two-tailed Student’s t-test). **(E)** Confocal images showing FLAG-miP-FERMT3 together with GM130 (Golgi apparatus), α-tubulin (microtubules), γ-tubulin (centrosome), ninein or CEP170 (centriole subdistal appendages) in endothelial cells. Nuclei were stained with DAPI (blue). Similar results were obtained in 3-5 independent cell batches. Scale bar = 25 µm.

### miP-FERMT3 interacts with proteins involved in protein catabolism and enhances ubiquitination and proteasome activity

MiPs often exert their biological function as components of macromolecular complexes [14–16]. To identify proteins that interact with miP-FERMT3, the FLAG-tagged fusion protein was expressed in endothelial cells, immunoprecipitated and the recovered complexes were subjected to mass spectrometry. This approach identified 129 miP-FERMT3-interacting proteins, most of which related to ubiquitin and proteasome-dependent protein catabolic processes (**Figure 2A-B, Dataset 1**). Among these were three non-ATPase regulatory subunits of the 26S proteasome. The top hit in the latter group was 26S proteasome non-ATPase regulatory subunit 9 (PSMD9), a ubiquitously expressed proteasomal chaperone that facilitates the assembly of the 26S proteasome and contributes to proteostasis in mammalian cells [17]. The interaction between miP-FERMT3 and PSMD9 was validated by co-immunoprecipitation and immunoblotting and was found to occur predominantly at the centrosome, as demonstrated by confocal microscopy (**Figure 2C-D**). Another miP-FERMT3-interacting protein involved in protein catabolism was cullin-2 (CUL2). This protein is a member of the Cullin–RING ligase complex, which is the largest family of RING-type E3 ubiquitin ligases and consists of a cullin scaffold, a RING-finger protein, an adaptor and a substrate-recognition module. This macromolecular complex mediates the ubiquitination of numerous target proteins. Confocal microscopy confirmed the colocalizaiton of miP-FERMT3 and CUL2 (**Figure 2E**). The activity of cullin is tightly regulated by post-translational modification with the ubiquitin-like molecule NEDD8, a process referred to as neddylation [18]. Notably, while NEDD8 appeared to be distributed throughout the nucleus in control cells, it co-localized with miP-FERMT3 in areas consistent with the centrosome in endothelial cells that expressed FLAG-miP-FERMT3 (**Figure 2F**). As our observations indicated that the miP could activate Cullin–RING ligases, we next determined whether protein ubiquitination was altered in miP-FERMT3-expressing cells. Global cellular ubiquitination was consistently increased in miP-FERMT3-expressing cells treated with MG132 to inhibit proteasomal degradation (**Fig 2G**). Notably, proteasomal activity, measured as degradation of a specific fluorogenic substrate, was also significantly enhanced in miP-FERMT3-expressing cells (**Figure 2H**).

**Figure 2.**
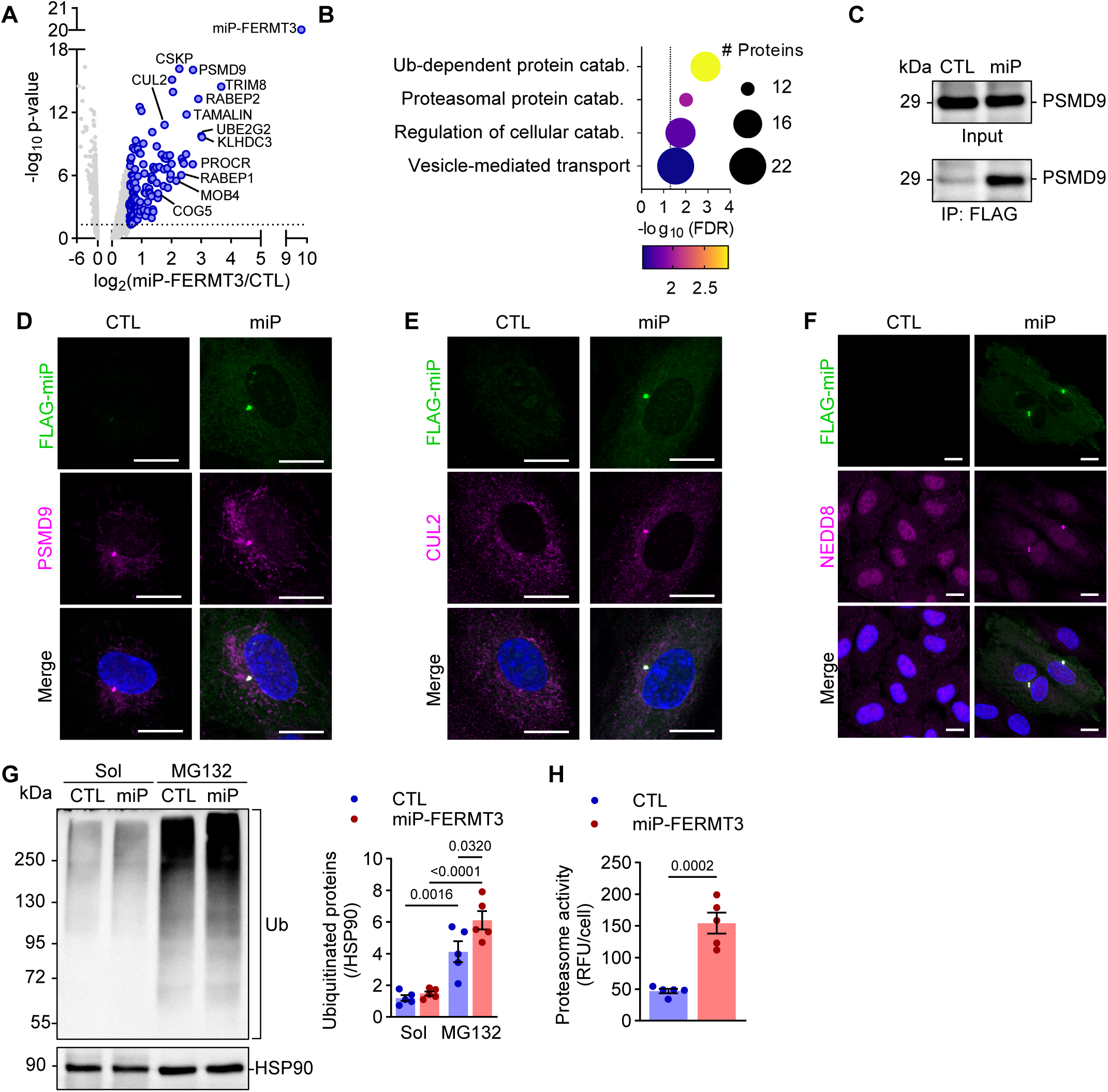
Interactome of miP-FERMT3 and effects on protein catabolism. **(A)** Volcano plot showing proteins associated with FLAG-miP-FERMT3; n=5 independent cell batches. The dashed line marks the significance threshold (p=0.05). **(B)** GO term enrichment analysis (STRING) of FLAG-miP-FERMT3 interacting proteins. The dashed line marks the significance threshold (FDR=0.05). **(C)** Co-immunoprecipitation (IP) of PSMD9 with FLAG-miP-FERMT3 from cell lysates (input) of endothelial cells expressing FLAG-miP-FERMT3 or EGFP (CTL). Similar results were observed in 3 independent cell batches. **(D-F)** Confocal images showing the colocalization of FLAG-miP-FERMT3 (FLAG-miP) with PSMD9 (D), CUL2 (E) and NEDD8 (F). Nuclei were stained with DAPI (blue). Similar results were obtained in 3 additional independent cell batches. Scale bars = 10 µm. **(G)** Protein ubiquitination (Ub) in endothelial cells expressing FLAG-miP-FERMT3 or EGFP (CTL) and treated with solvent (Sol) or MG132 (10 μmol/L,16 hours); n=5 independent cell batches (two-way ANOVA and Tukey’s multiple comparison). **(H)** Proteasomal activity in endothelial cells expressing FLAG-miP-FERMT3 or EGFP (CTL); n=5 independent cell batches (unpaired, two-tailed Student’s t-test).

### miP-FERMT3 induces nuclear accumulation of p53 and cell cycle arrest

TRIM8, a member of the RING-type E3 ubiquitin ligase family was one of the proteins that associated with miP-FERMT3, an interaction that was validated by co-immunoprecipitation and immunoblotting (**Figure 3A**). TRIM8 regulates a broad range of cellular processes [19] as well as being a tumor suppressor by virtue of its ability to stabilize and activate p53. In turn, p53 transcriptionally induces TRIM8 expression, thereby establishing a positive feedback loop that reinforces p53 activity to induce cell cycle arrest [20]. Consistent with our observation that miP-FERMT3 interacted with TRIM8, the expression of miP-FERMT3 resulted in marked nuclear accumulation of p53, with approximately 75% of miP-FERMT3-expressing endothelial cells exhibiting nuclear p53 positivity (**Figure 3B**). TRIM8 protein levels were also significantly increased, indicating that a TRIM8-p53 regulatory loop is active in the presence of miP-FERMT3 (**Figure 3C**).

**Figure 3.**
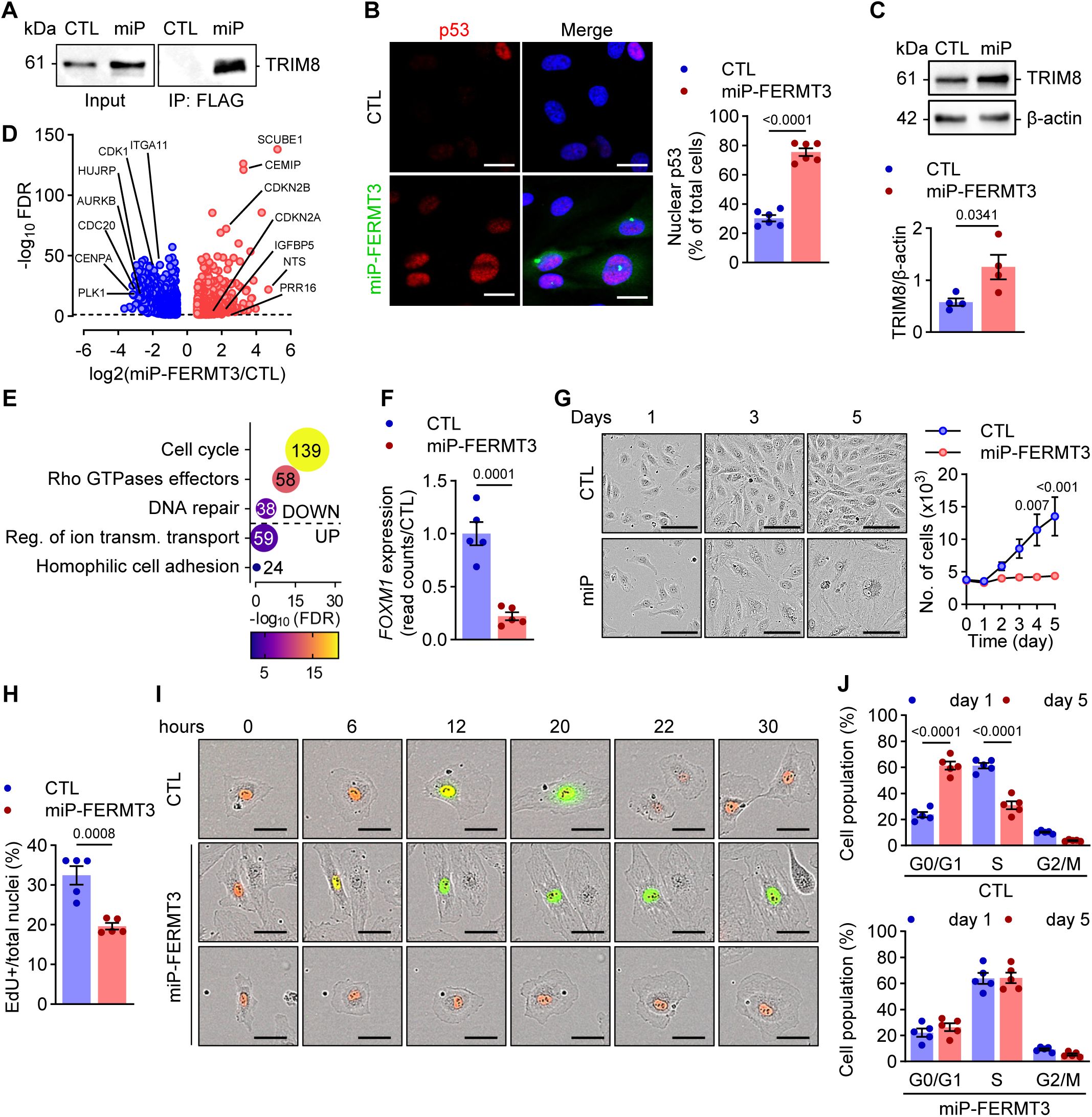
Impact of miP-FERMT3 expression on the nuclear accumulation of p53 and cell-cycle regulation. **(A)** Co-immunoprecipitation (IP) of TRIM8 and FLAG-miP-FERMT3 from endothelial cell lysates (input) expressing FLAG-miP-FERMT3 or EGFP (CTL). Similar results were observed in 4 independent cell batches. **(B)** Confocal images showing p53 in endothelial cells expressing FLAG-miP-FERMT3 or EGFP (CTL); n=6 independent cell batches (unpaired, two-tailed Student’s t-test). Scale bar = 25 µm. **(C)** Levels of TRIM8 in cells expressing FLAG-miP-FERMT3 or EGFP (CTL); n=4 independent cell batches (unpaired, two-tailed Student’s t-test). **(D-E)** Volcano plot (D) and gene set enrichment analysis (STRING with ranking – Reactome) (E) of upregulated (red) and downregulated (blue) genes in endothelial cells expressing FLAG-miP-FERMT3 or FLAG-miRFP670nano (CTL). Numbers inside bubble plot indicate # of genes/GO term; n=5 independent cell batches. The dash line in D marks the significance threshold (FDR=0.05). **(F)** RNA expression (read counts) of *FOXM1* in endothelial cells expressing FLAG-miP-FERMT3 or miRFP647nano (CTL); n=5 independent cell batches (unpaired, two-tailed Student’s t-test). **(G)** Representative images of growth factor-induced proliferation of endothelial cells expressing FLAG-miP-FERMT3 or EGFP (CTL) and respective quantification of number of cells per well; n=5 independent cell batches (two-way ANOVA and Šidák’s multiple comparison). Scale bar: 100 µm. **(H)** EdU incorporation in cells expressing FLAG-miP-FERMT3 or EGFP (CTL); n=5 independent cell batches (unpaired, two-tailed Student’s t-test). **(I)** Time course (up to 30 hours) of changes in FUCCI fluorescence in endothelial cells expressing FLAG-miP-FERMT3 or empty vector (CTL) . Red: G1 phase, yellow: G1/S, green: S/G2/M. Similar results were obtained in 5 independent cell batches. Scale bar: 50 µm. **(J)** Cell cycle progression (flow cytometry) of cells expressing FLAG-miP-FERMT3 or EGFP (CTL) showing DNA content distribution at day 1 and day 5 post-transduction; n=5 independent cell batches (two-way ANOVA and Šídák’s multiple comparisons test).

The actions of p53 have been extensively characterized and it regulates the expression of genes involved in DNA damage, cell cycle progression, apoptosis and senescence [21, 22]. To determine the impact of miP-FERMT3 on transcription, we performed whole-transcriptome RNA sequencing of endothelial cells lacking or expressing the FLAG-miP-FERMT3. This analysis identified a total of 1410 miP-FERMT3-regulated genes, 639 of which were upregulated and 771 downregulated. Genes down-regulated by the miP included a series of genes related to cell cycle progression, including cyclin-dependent kinase 1 (*CDK1*), polo-like-kinase 1 (*PLK1*), aurora kinase B (*AURKB*), and DNA repair genes such as radiation sensitive protein 51 (*RAD51*), fanconi anemia group D2 (*FANCD2*), and exonuclease 1 (*EXO1*). In contrast, genes associated with cell cycle inhibition and senescence, including the cyclin-dependent kinase inhibitors *CDKN2A* and *CDKN2B*, were markedly upregulated in miP-expressing cells (**Figure 3D-E**, **Dataset 2**). Despite the fact that miP-FERMT3 increased cellular/nuclear p53 levels it did not induce the expression of classical p53-regulated genes such as *CDKN1A* (p21), *GADD45A* and *CCNG1*. Next, transcription factor enrichment analysis using all differentially expressed genes (ChEA3 platform [23]) predicted that forkhead box M1 (*FOXM1*) was the transcription factor most likely to be responsible for the downregulation of a large proportion of the genes suppressed in miP-FERMT3-expressing endothelial cells (**Dataset 3**). This is of particular relevance inasmuch as p53 is a well-established negative regulator of FOXM1 [24, 25], which was also strongly downregulated in miP-FERMT3-expressing endothelial cells (**Figure 3F**).

In line with the observed transcriptional alterations, cell growth was abrogated in miP-FERMT3-expressing endothelial cells (**Figure 3G**). Consistent with this, EdU incorporation was significantly reduced by miP-FERMT3 expression (**Figure 3H**), indicating that fewer cells progressed into the S/G2/M phases. When the fluorescent ubiquitination-based cell cycle indicator (FUCCI) was used to visualise cycling cells, we observed that miP-FERMT3 induced early cell cycle defects and cell cycle arrest at both the G1 (red) and S/G2/M (green) phases (**Figure 3I**). Flow cytometry analyses of DNA content using Hoechst staining confirmed pronounced cell cycle arrest in miP-FERMT3-expressing cells (**Figure 3J**). Even after five days of growth factor stimulation, miP-FERMT3-expressing cells remained in growth arrest with the majority (∼60%) of cells in S phase, followed by G1 (∼20%) and G2/M (∼10%).

As cells progress through the cell cycle, centrosomes are duplicated and mature structurally to ensure proper mitosis. This process involves the expansion of the pericentriolar matrix and the sequential assembly of distal and subdistal appendage proteins at centrioles [26]. The centriole subdistal appendage protein ninein initially localizes to the mother centriole and gradually distributes to the daughter centriole as centrosome duplication advances [27]. Another component of the centrosome, CEP170, is essential for recruiting and stabilizing microtubules at the subdistal appendages [28]. We observed an accumulation of both ninein and CEP170 in miP-FERMT3 expressing endothelial cells (**Figure 4A-D**). Similarly, levels of the cell cycle-dependent CDK substrate CP110, a key regulator of centrosome duplication, were also elevated in the presence of miP-FERMT3 (**Figure 4E**).

**Figure 4.**
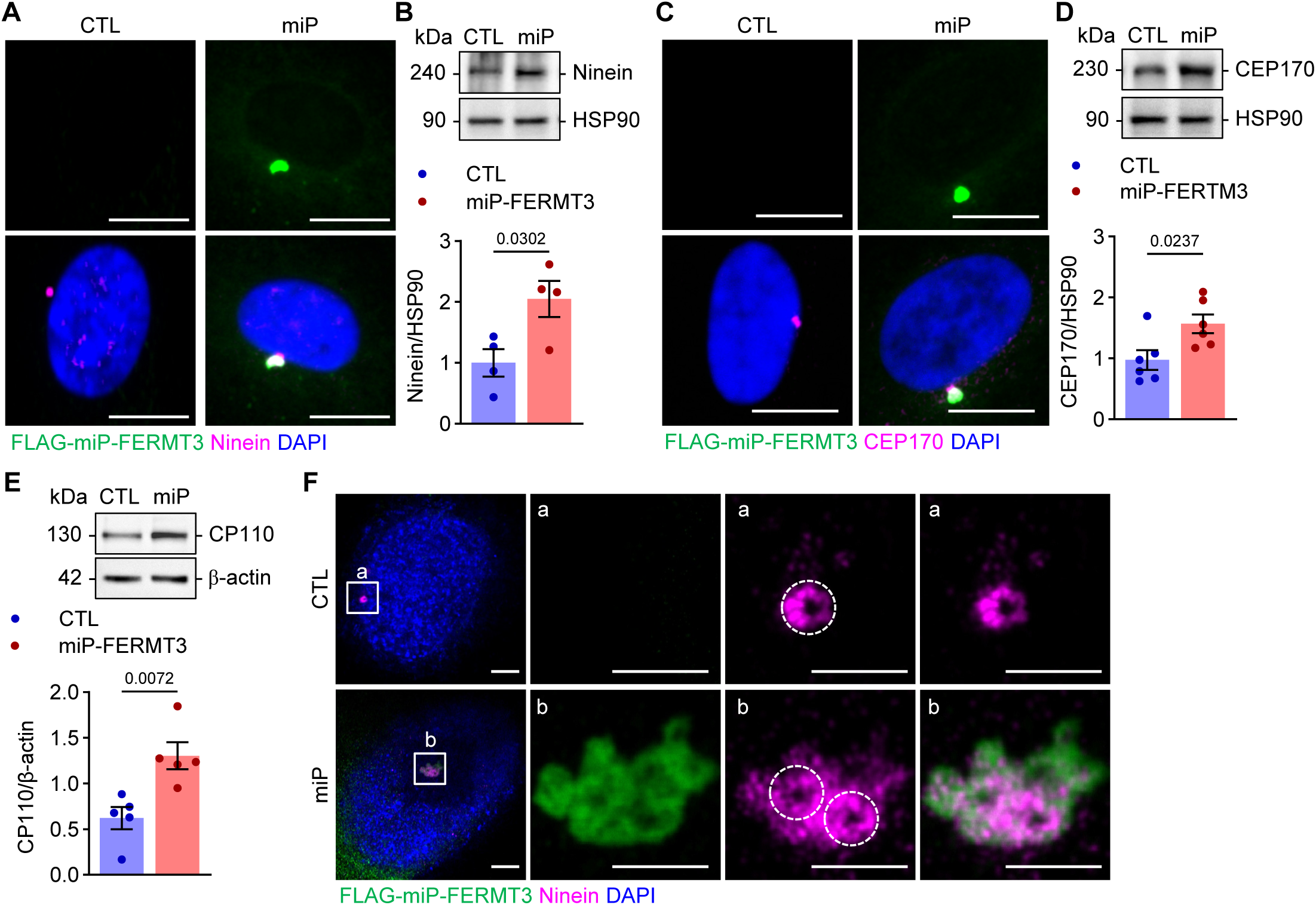
Consequence of miP-FERMT3 expression on centrosomal proteins. **(A-B)** Representative confocal microscopy images (A) and levels (B) of ninein in endothelial cells overexpressing FLAG-miP-FERMT3 or EGFP (CTL). Similar results were obtained in 4-5 independent cell batches (unpaired, two-tailed Student’s t-test). Scale bar: 10 μm. **(C-D)** Representative confocal microscopy images (C) and levels (D) of CEP170 in endothelial cells overexpressing FLAG-miP-FERMT3 or EGFP (CTL). Similar results were obtained in 4-6 independent cell batches (unpaired, two-tailed Student’s t-test). Scale bar: 10 μm. **(E)** Levels of CP110 in endothelial cells overexpressing FLAG-miP-FERMT3 or EGFP (CTL); n=5 independent cell batches (unpaired, two-tailed Student’s t-test). **(F)** Representative expansion microscopy images showing coiled-coil images of ninein in endothelial cells overexpressing FLAG-miP-FERMT3 or EGFP (CTL). The regions highlighted as a and b are shown at higher magnification next to the first image. Similar results were obtained in 5 independent cell batches. Scale bars: 10 µm and 5 µm (a and b).

To better visualize structural changes in the subdistal appendages of centrioles we used expansion microscopy. In control endothelial cells, ninein clustered into a ring, which is known to surround the centriole [29]. In the cell cycle arrested miP-FERMT3-expressing cells ninein clusters formed multiple interconnected ring-like structures, indicative of centrosome duplication or amplification (**Figure 4F**). These observations suggest that miP-FERMT3-expressing cells accumulate duplicated centrosomes as a consequence of incomplete mitotic progression.

### Overexpression of miP-FERMT3 induces cellular senescence

Replicative stress and the resulting dysregulation of centrosomal proteins contribute to genomic instability [30]. Because genes involved in DNA repair were supressed in miP-FERMT3-expressing endothelial cells, we next looked for evidence of DNA damage and found γ-H2AX foci in miP expressing cells (**Figure 5A**). Live cell imaging further revealed that already 2 days after transduction, miP-FERMT3-expressing cells were markedly larger than cells transduced using a control adenovirus, a phenomenon that became more pronounced with time (**Figure 5B**).

**Figure 5.**
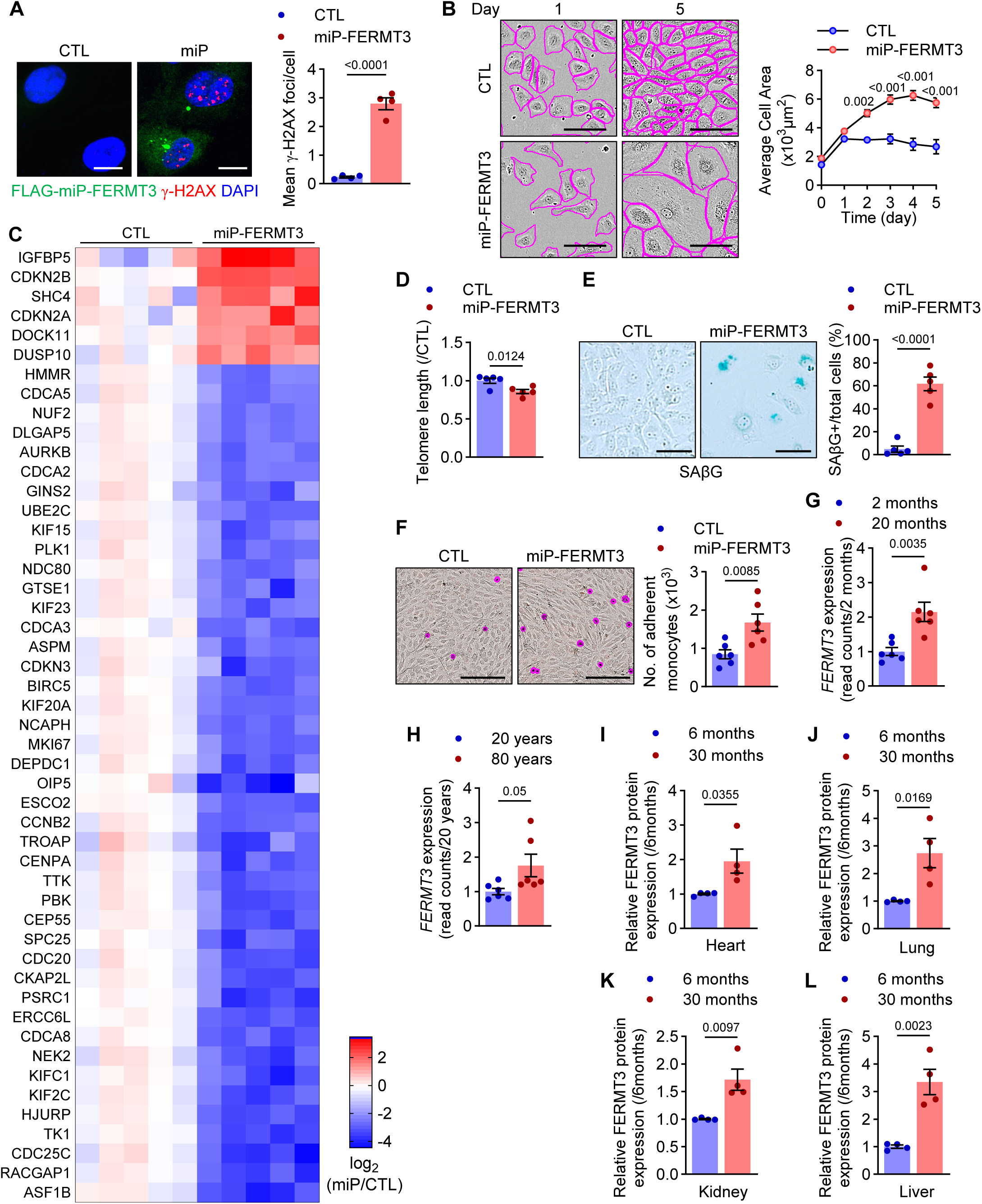
Impact of miP-FERMT3 on cellular senescence. **(A)** γ-H2AX foci in endothelial cells expressing FLAG-miP-FERMT3 or EGFP (CTL). n=4 independent cell batches (unpaired, two-tailed Student’s t-test). Scale bar: 10 µm. **(B)** Cell size in cells expressing FLAG-miP-FERMT3 or EGFP (CTL); n=6 independent cell batches (two-way ANOVA and Šidák’s multiple comparison). Scale bar: 100 µm. **(C)** Senescence-associated genes significantly (FDR≤0.05) altered by FLAG-miP-FERMT3 versus FLAG-miRFP670nano (CTL) expression; n=5 independent cell batches. **(D)** Relative changes in telomere length in cells expressing FLAG-miP-FERMT3 or EGFP (CTL); n=5 independent cell batches (unpaired, two-tailed Student’s t-test). **(E)** SAβG in cells expressing FLAG-miP-FERMT3 or EGFP (CTL); n=6 independent cell batches (unpaired, two-tailed Student’s t-test). Scale bar: 100 µm. **(F)** Adhesion of monocytes (magenta) to endothelial cell monolayers following incubation with conditioned medium from cells expressing FLAG-miP-FERMT3 or EGFP (CTL); n=6 independent cell batches (unpaired, two-tailed Student’s t-test). Scale bar: 200 µm. **(G)** *FERMT3* transcript expression (read counts) in cardiac endothelial cells from 2-versus 20-month old mice; n=6 mice/group (unpaired, two-tailed Student’s t-test). **(H)** *FERMT3* transcript expression (read counts) in mesenteric artery endothelial cells from 20-versus 80-year old subjects; n=6 individuals/group (unpaired, two-tailed Student’s t-test). **(I-L)** FERMT3 protein levels in hearts (I), lungs (J), kidneys (H) and livers (L) from 6-versus 30-month old mice; n=4 mice/group (unpaired, two-tailed Student’s t-test).

Persistent cell cycle arrest can lead to replicative senescence and the appearance of “giant cells” frequently mirrors the development of cellular senescence [31]. We therefore intersected the differentially regulated genes in miP-FERMT3-expressing endothelial cells with a curated dataset of senescence-associated genes using Senescent Cell Identification (SenCID) [32]. This analysis revealed a robust senescence-associated gene expression signature in miP-FERMT3-expressing cells (**Figure 5C**). Further validation of the induction of senescence was provided by estimating telomere length and senescence-associated β-galactosidase (SAβG) activity. Indeed, miP-FERMT3 expression resulted in a significant shortening of telomeres, as quantified by monochrome multiplex qPCR (**Figure 5D**). Moreover, approximately 60% of the miP-expressing cells expressed senescence-associated β-galactosidase (SAβG) within 7 days of transduction (**Figure 5E**). This is a significant finding as repetitive passaging of cells seeded at low density for up to 10 times is generally required for human endothelial cells to express SAβG [33] and the cells studied were passaged maximally six times. Indeed, SAβG was largely undetectable in the cells transduced with the control virus. Cellular senescence has been associated with a change in the secretome that includes the release of pro-inflammatory mediators [34, 35]. In line with this, the application of medium from miP-FERMT3-overexpressing cells to untreated endothelial cells induced inflammatory cell activation as evidenced increased monocyte adhesion compared with cells treated with culture medium from miP-deficient cells (**Figure 5F**).

The vasculature is among the first systems to “age” and has even been proposed to orchestrate systemic aging [10, 36]. To determine whether the expression of FERMT3, which hosts miP-FERMT3, is affected by aging we interrogated publicly available datasets that compared endothelial cell transcriptomes from young and aged mice and humans [33, 37]. This revealed higher *FERMT3* expression in endothelial cells from 20 month old versus 2 month old mice (**Figure 5G**). Similarly, endothelial cells from 80 year old humans expressed higher *FERMT3* expression than those from 20 year old subjects (**Figure 5H**). Data from a recent aging mouse proteome atlas [38], confirmed these findings as FERMT3 protein was significantly more abundant in the heart, lungs, liver and kidneys of 30 month versus 6 month old mice (**Figure 5I-L**).

## DISCUSSION

In this study, we characterized the function of miP-FERMT3, a previously unannotated miP encoded within the *FERMT3* gene in human endothelial cells. Functionally, miP-FERMT3 expression could be linked to cell cycle arrest, centrosome amplification and DNA damage, leading to rapid endothelial senescence and a paracrine pro-inflammatory phenotype. Notably, *FERMT3* expression was elevated in aged mouse and human endothelial cells, suggesting a conserved role of miP-FERMT3 in vascular senescence and aging. At the molecular level miP-FERMT3 associated with centriole subdistal appendages, where it interacted with components of the protein catabolic machinery, enhancing neddylation, ubiquitination, and proteasomal activity. miP-FERMT3 also stabilized p53, repressing cell cycle and DNA repair genes while inducing senescence-associated transcriptional programs.

The location of a miP in a cell frequently gives hints about its likely cellular function. Indeed, miPs that associate with membrane receptors modulate downstream signal transduction pathways, while miPs that regulate cell metabolism have been localized to mitochondria [39–41]. In endothelial cells, FLAG-miP-FERMT3 was detected diffusely distributed through the cytosol as well as in distinct hotspots close to the nucleus. The latter were identified as centrosomes by virtue of colocalization with the centrosomal markers α- and γ-tubulin, as well as the centriole subdistal appendage protein markers ninein and CEP170.

Centrosomes play an essential role in the control of cell cycle progression, cytoskeletal organization, the DNA damage response and proteostasis [11, 13, 39]. Indeed, miP-FERMT3 in centrosomes physically interacted with multiple proteins implicated in ubiquitin- and proteasome-dependent protein catabolism. Among these were PSMD9, a 19S regulatory subunit of the 26S proteasome complex [42], CUL2, an E3 ubiquitin ligase complex scaffold protein [43] and TRIM8, an E3 ubiquitin ligase [44]. The ubiquitination cascade is orchestrated by a series of enzymes: E1 ubiquitin-activating, E2 ubiquitin-conjugating and E3 ubiquitin-ligating enzymes, along with deubiquitinases that remove ubiquitin moieties. RING-finger E3 ligases represent the largest and most diverse family among the E3 ligases and include the cullin-RING E3 ligase family [45], all of which are regulated by post-translation modification with NEDD8 [18]. Importantly, NEDD8 accumulated in centrosomes in miP-FERMT3-expressing cells implying the activation of cullin-RING E3 ligases. Finding miP-FERMT3 together with PSMD9 and CUL2 implied a role of the miP in regulating E3 ligase function and protein ubiquitination. Although the impact of miP-FERMT3 on global cellular protein ubiquitination was modest, it did significantly increase proteasome activity. The interaction of miP-FERMT3 with TRIM8 was particularly interesting as it is a RING-type E3 ubiquitin ligase that acts as a tumour suppressor by stabilizing and activating p53, thereby promoting cell cycle arrest. It does this by preventing the interaction of p53 with MDM2 [20] . TRIM8 and p53 are also part of a positive feedback loop as p53 induces TRIM8 expression [44, 46]. Indeed, miP-FERMT3 expression not only resulted in the nuclear accumulation of p53, it also increased TRIM8 expression.

The activation of p53 induces G1- or G2/M phase arrest, allowing DNA repair and preventing propagation of damaged DNA [47]. Consistent with its effects on p53, the introduction of miP-FERMT3 into endothelial cells decreased the expression of cell cycle-related genes and abrogated proliferation. However, the genes regulated by miP-FERMT3 were not the expected classical p53 target genes. Instead, it was possible to identify FOXM1 as underlying the changes observed. FOXM1 is a key regulator of cell cycle progression, driving the expression of genes required for DNA replication, mitosis and genomic stability [24, 25] and is negatively regulated by p53 [22, 23]. A decrease in FOXM1 expression is also a hallmark of cellular senescence, leading to irreversible cell cycle arrest and centrosome amplification through impaired/incomplete transitions through G1/S and G2/M [48–51]. During S-phase, the centrosome duplicates and expands its pericentriolar matrix while assembling distal and subdistal appendage proteins such as ninein and CEP170 [26]. Ninein also functions as a component of centrosome linkers which maintains centriole cohesion until its dissolution in late G2 [52, 53]. The excessive accumulation of ninein aggregates, as well as of CEP170, and CP110 in miP-FERMT3-expressing cells indicates that centrosome duplication was initiated but not successfully completed, consistent with a failure to progress through the G2/M transition, explaining why miP-expressing cells failed to proliferate. Such centrosome duplication defects are known triggers of a robust senescence response [54]. Consequently miP-FERMT3 expression resulted in rapid phenotypic changes consistent with cellular senescence, including increased cell size, cell cycle arrest, the expression of SAβG, loss of proteostasis, telomere attrition, DNA damage and the secretion of factors that promoted endothelial activation and increased monocyte adhesion to the cell surface. Given the strong link between vascular senescence and aging, we sought to determine whether expression of miP-FERMT3 was altered in aging *in vivo*. Direct assessment of miP-FERMT3 levels was not feasible due to the lack of tools to detect the endogenous miP. Therefore, we used *FERMT3* transcript and protein abundance as a proxy for miP-FERMT3 expression. Analysis of human and murine endothelial datasets revealed that *FERMT3* is upregulated with age, supporting a potential role for miP-FERMT3 in driving vascular senescence and age-associated endothelial dysfunction.

Taken together, this study demonstrates that miP-FERMT3 modulates the ubiquitin-proteasome system and elicits p53-dependent cell cycle arrest and cellular senescence. The upregulation of the miP in the context of inflammation may contribute to the onset of vascular senescence that is a hallmark of aging.

## METHODS

### Cell culture

Human umbilical vein endothelial cells were isolated and cultured as described previously [55, 56] and used between passage 4-6. The use of human material in this study complies with the principles outlined in the Declaration of Helsinki (World Medical Association, 2013), and the isolation of endothelial cells was approved in written form by the ethics committee of the Goethe-University. THP-1 monocytic cells were obtained from the American Type Culture Collection (LGC Standards; Hamburg, Germany) and cultured in RPMI-1640 containing 2 mmol/L glutamine, 10 mmol/L HEPES, 1 mmol/L sodium pyruvate, 4.5g/L glucose, 1.5 g/L sodium bicarbonate and 10% foetal calf serum. AD-293 cells were cultured as described previously [57]. All cell cultures tested negative for mycoplasma contamination. Cells were maintained in a humidified incubator at 37 °C with 5% CO₂.

To assess K48-linked polyubiquitination, cells were treated with 10 μmol/L MG132 (Cat. # M7449; Merck; Darmstadt, Germany) or 0.1% dimethyl sulfoxide (DMSO) as solvent control in endothelial cell growth medium 2 (ECGM2; PromoCell, Heidelberg, Germany) for 16 hours at 37 °C.

### Generation of adenoviruses, lentiviruses and cell transduction

#### Adenovirus generation and transduction

The adenoviral vector used to overexpress FLAG-tagged miP-FERMT3 was constructed and packaged by VectorBuilder. Briefly, the sequence encoding the FLAG tag (GACTACAAAGACGATGACGACAAG) was inserted at the 5’ end of the human smORF-FERMT3 sequence (GCCGGCCCCAGCCGCCACCCGACCCCCTCCTGCTCCAGCGTCTGCCACGGCCCAGCTCCCTGTCAGACAAGACCCAGCTCCACAGCAGGTGGCTGGACTCGTCGCGGTGTCTCATGCAGCAGGGCATCAAGGCCGGGGACGCACTCTGGCTGCGCTTCAAGTACTACAGCTTCTTCGATTTGGATCCCAAGACAGACCCCGTGCGGCTGA), preceded by Kozak sequence (GCCACC) and ATG start codon. The restriction sites AbsI and SgrDI were placed on the 5’ and 3’ end on of the insert, respectively (VectorBuilder ID: VB230707-1073jqc). The insert was cloned into the mammalian gene expression adenoviral vector pAd5 under the cytomegalovirus (CMV) promoter. DNA sequencing confirmed correct direction of the insert and lack of mutations. The adenoviral vector and replication incompetent adenoviruses used to overexpress FLAG-tagged miRFP670nano or EGFP were also constructed and packaged by VectorBuilder as described above (VectorBuilder ID: VB230223-1677bmd, VB010000-9299hac). FLAG-miRFP670nano was used as control as it represents smallest near infra-red fluorescent protein. Alternatively, for experiments involving the Fluorescent Ubiquitination-based Cell Cycle Indicator, adenoviruses carrying an empty pAdShuttle-CMV vector were generated as described [58]. Generation and expansion of all replication incompetent adenoviruses was carried out by transfection of the packaging cell line AD-293. Human endothelial cells (passage 4, 90% confluency) were starved of serum in endothelial cell basal medium (EBM, Cat. # C-22211; PromoCell, Heidelberg, Germany) containing 0.1% bovine serum albumin (Cat. # A8412; Sigma-Aldrich; Darmstadt, Germany) and infected with adenoviruses (250 MOI) overnight. On the following day, the medium was replaced with ECGM2 supplemented with 10% foetal calf serum (Sigma-Aldrich; Darmstadt, Germany).

#### Lentivirus generation and transduction

Lentiviruses to express the Fluorescent Ubiquitination-based Cell Cycle Indicator were generated as described [59]. Human endothelial cells were transduced with two lentiviruses to express mCherry-hCdt1 and mAG-hGeminin for 24 hours in the presence of 10 µg/mL polybrene (Cat. # sc-134220; Santa Cruz Biotechnology, Heidelberg, Germany). Following transduction, cells were cultured for an additional 24 hours before further experiments.

### Immunofluorescence

Cells grown on 8 well chamber slides (Ibidi, Martinsried, Germany) were washed with phosphate buffered saline (PBS) containing 0.8 mmol/L CaCl_2_ and 1.4 mmol/L MgCl_2_. For mitochondrial staining, cells were incubated with 100nmol/L MitoTracker® Red CMXRos (Cat. # M7512, ThermoFisher Scientific; Darmstadt, Germany) for 30 minutes in a humidified incubator at 37°C. Then cells were fixed in ice cold methanol (Sigma-Aldrich; Darmstadt, Germany) or 4% paraformaldehyde (ThermoFisher Scientific; Darmstadt, Germany) and incubated in blocking/permeabilization buffer containing 5% horse serum and 0.1% Triton X-100 (Cat. 3051.2, Carl Roth, Karlsruhe, Germany) in PBS for 30 minutes at room temperature, followed by incubation with anti-calnexin (Cat. # C4731, Sigma-Aldrich; Darmstadt, Germany), anti-CEP170 (Cat. # 27325-1-AP, Proteintech; Planegg-Martinsried, Germany), anti-CUL2 (Cat. # 67175-1-IG, ThermoFisher Scientific, Darmstadt, Germany), anti-EEA1 (Cat. # 3288, Cell Signaling Technology; Leiden, Netherlands), anti-FLAG (Cat. # F3165 or # F7425, Sigma-Aldrich; Darmstadt, Germany), anti-GM130 (Cat. # ab52649, Abcam; Amsterdam, Netherlands), anti-LAMP1 (Cat. # 9091, Cell Signaling Technology; Leiden, Netherlands), anti-NEDD8 (Cat. # ab81264, Abcam; Amsterdam, Netherlands), anti-ninein (Cat. # ab4447, Abcam; Amsterdam, Netherlands), anti-NOGO-B (Cat. # ab47085, Abcam; Amsterdam, Netherlands), anti-p53 (Cat. # 2527S; Cell Signaling Technology; Leiden, Netherlands), anti-PSMD9 (Cat. # ab233154, Abcam; Amsterdam, Netherlands), anti-α-tubulin (Cat. # 2125, Cell Signaling Technology; Leiden, Netherlands), anti-γ-H2AX (Cat. # ab81299, Abcam; Amsterdam, Netherlands), anti-γ-tubulin (Cat. # ab11317, Abcam; Amsterdam, Netherlands) in 0.5% horse serum and 0.01% Triton X-100 in PBS for 2 hours at room temperature. Cells were then incubated with donkey anti-mouse Alexa Fluor 488 (Cat. # A-21202; Invitrogen, Darmstadt, Germany) and donkey anti-rabbit Alexa Fluor 555 (Cat. # A-32794; Invitrogen, Darmstadt, Germany) in PBS for 1 hour at room temperature. Nuclei were stained using 4′,6-diamidino-2-phenylindole (DAPI) (Cat. # A1001, Applichem GmbH; Darmstadt, Germany). Finally, cells were overlaid with mounting medium containing 50% (v/v) glycerol (Cat. # 3783.1, Carl Roth, Karlsruhe, Germany) and dithiothreitol (DTT, 1 mol/L) (Cat. # A2948, Applichem GmbH; Darmstadt, Germany). Images were taken using an SP8 (Leica, Wetzlar, Germany) or LSM-780 confocal microscope (Zeiss, Jena, Germany) and LAS AF lite software (Leica) or ZEN software (Zeiss).

The mean fluorescence intensity (MFI) of p53 and mean γ-H2AX foci were quantified using ImageJ (version 1.54d) software. MFI values for p53 were normalized to the MFI of DAPI to account for nuclear area, and results were expressed as the percentage of total nuclear p53. The number of p53 positive cells that were FLAG-miP-positive or FLAG-miP-negative were quantified and expressed as percentage of p53 cells. For each experimental cell batch, 3-5 different fields were imaged and analyzed for quantification. For quantification of γ-H2AX foci, at least 200 cells per batch were analyzed, and the mean number of γ-H2AX foci per cell was calculated.

### Expansion microscopy

Preparation for expansion microscopy was done as described previously [60, 61]. Briefly, cultured cells for expansion microscopy were fixed on 13 mm coverslips ice cold methanol for 15 minutes. Cells were immunostained with anti-Ninein (Cat. # ab4447; Abcam; Amsterdam, Netherlands), and anti-FLAG (Cat. # F3165; Sigma-Aldrich; Darmstadt, Germany) antibodies for 3 days and then donkey anti-mouse Alexa Fluor 488 (Cat. # A-21202; Invitrogen, Darmstadt, Germany), donkey anti-rabbit Alexa Fluor 555 (Cat. # A-32794; Invitrogen, Darmstadt, Germany) and DAPI in PBS for 2 days. Stained cells on coverslips were transferred to 35 mm MatTek dishes (P35G-1.0-14-C, MatTek) and incubated with 10 mg/ml Acryloyl-X/DMSO in PBS (1:100) for 3 hours at room temperature. Samples were washed twice with PBS for 15 minutes. Samples were subsequently incubated with a freshly prepared gelling solution composed of Stock X, tetramethylethylenediamine, ammonium persulfate, and distilled water in a 47:1:1:1 (v/v/v/v) ratio. Stock X contained 0.914 mol/L sodium acrylate, 0.352 mol/L acrylamide, 9.7 mmol/L N,N′-methylenebisacrylamide, 2 mol/L NaCl, and 1× PBS in water. Then, samples were covered with a glass 15mm coverslip and incubated at 37°C for 2 hours. After the gel polymerized, the coverslips were removed and the gel was incubated with an aqueous digestion buffer containing 0.5% Triton X-100, 1 mmol/L EDTA disodium pH 8.0, 50 mmol/L Tris-HCl pH 8.0, 800 mmol/L NaCl and 8 U/ml Proteinase K on a shaker overnight at room temperature. After removal of the digestion buffer, gel samples were incubated with distilled water 4 times for 20 minutes. Samples were embedded in 4% low-melting point agarose in water in a 35 mm dish containing a polymer coverslip (81151, Ibidi). Images were taken using a confocal microscope (LSM-780; Zeiss, Jena, Germany) and ZEN software (Zeiss).

### Cell lysis and immunoblotting

Samples were lysed in ice-cold radioimmunoprecipitation assay buffer (50 mmol/L Tris HCl pH 7.5, 150 mmol/L NaCl, 25 mmol/L NaF, 10 mmol/L Na_4_P_2_O_7_, 1% Triton X-100 and 0.5% sodium deoxycholate) supplemented with 0.1% sodium dodecyl sulfate (SDS) and protease and phosphatase inhibitors. Protein concentrations were determined using the Bradford assay, and detergent-soluble proteins were solubilized in sample buffer containing 2% SDS, 1% β-mercaptoethanol and 0.005% bromophenol, separated by SDS-PAGE and subjected to immunoblotting. Membranes were incubated with anti-ninein (AB4447, Abcam; Amsterdam, Netherlands), anti-CEP170 (Cat. # 27325-1-AP, Proteintech; Planegg-Martinsried, Germany), anti-TRIM8 (Cat. # ab316149, Abcam; Amsterdam, Netherlands), anti-K48 polyubiquitin (Cat. # 8081, Cell Signaling Technology; Leiden, Netherlands), anti-CP110 (Cat. # 12780-1-AP, ThermoFisher Scientific; Darmstadt, Germany), anti-PSMD9 (Cat. # PA5121663, ThermoFisher Scientific; Darmstadt, Germany), anti-HSP90 (Cat. # 610419, BD Biosciences; Heidelberg, Germany), anti-β-actin (Cat. # MA1115, Boster Biologics; Hamburg, Germany). Proteins were visualized by enhanced chemiluminescence using a commercially available kit SuperSignal™ West Femto Maximum Sensitivity Substrate (Cat. # 34095; ThermoFisher Scientific; Darmstadt, Germany).

### Immunoprecipitation

Three days after adenoviral transduction, cells were washed once with PBS prior to collection in 700 µl of affinity purification lysis buffer containing Tris/HCl pH 7.5 (50 mmol/L), NaCl (150 mmol/L), NP-40 (1%), Na_4_P_2_O_7_ (10 mmol/L), NaF (20 mmol/L), orthovanadate (2 mmol/L), okadaic acid (10 nmol/L), β-glycerophosphate (50 mmol/L), phenylmethylsulfonyl fluoride (230 µmol/L) and an EDTA-free protease inhibitor mix (Applichem GmbH; Darmstadt, Germany GmbH, Darmstadt, Germany). Lysates were incubated for 45-60 minutes on an end-over-end rocker and gently vortexed until no cell clumps were visible. Protein concentration was determined using the Bradford method. Whole cell lysates (500µg/sample) were incubated with 20µl anti-FLAG affinity gel (A2220, Merck; Darmstadt, Germany) overnight on an end-over-end rocker. Samples were then centrifuged at 5000 rpm for 2 minutes and the supernatant was removed. FLAG immunoprecipitates were washed twice with affinity purification lysis buffer and centrifuged at 5000 rpm for 30 seconds at 4^°^C. For proteomic analysis, FLAG immunoprecipitates were washed with a buffer containing 50 mmol/L Tris HCl (pH 7.5) and 150 mmol/L NaCl, heated in elution buffer containing Tris HCl pH 7.5 (50 mmol/L) and 2% SDS for 10 minutes at 95^°^C and stored at -20°C until further processing. For immunoblotting, FLAG immunoprecipitates were directly heated in elution buffer containing 2% SDS, 1% β-mercaptoethanol and 0.005% bromophenol blue in PBS for 10 minutes at 95^°^C.

### Identification of the miP-FERMT3 interactome by LC-MS/MS

miP-FERMT3 interactomes were prepared for mass spectrometry-based proteomics using a modified single-pot, solid-phase-enhanced sample preparation (SP3) method [62], adapted for a 96-well plate format. As proteins were eluted in an SDS-containing buffer, the addition of a detergent for initial denaturation was not required. Sample preparation was initiated immediately with the thermal denaturation and reduction steps. Samples were incubated for 10 minutes at 90°C with shaking at 1200 rpm in a ThermoMixer to ensure full denaturation, followed by the addition of the DTT reductant to begin the reduction step. The remainder of the SP3 protocol, including the subsequent alkylation, bead binding, extensive washing (to remove the SDS detergent), tryptic digestion and final peptide elution was performed as described on zenodo.org, record number: 17474920.

Peptide injection, separation and measurement on Bruker TimsTof Ultra as well as subsequent spectrum matching with DIA-NN [63] was performed as described on zenodo.org, record number: 17475274. The search was performed against the Human Uniprot.org database (reviewed Swiss-Prot entries; October 2022) complemented with the FLAG-miP sequence used (database is available in the repository, see section Data and materials availability).

Downstream data processing and statistical analysis were carried out by the Autonomics package developed in-house (10.18129/B9.bioc.autonomics:1.15.235). Proteins with a q-value of <0.01 were included for further analysis. MaxLFQ log2maxlfq intensities were used for quantitation and missing values imputed. All intensities containing only 1 precursor (Np) per sample were exchanged by NA for that particular sample. Differential abundance of protein groups was evaluated by limma [64].

The full list of DIA-NN settings, DIA-NN output, R code for data processing and statistical analysis are uploaded along with the mass spectrometric raw data to the ProteomeXchange Consortium (see section Data and materials availability).

For the miP-FERMT3 interactome, proteins significantly (FDR≤0.05) enriched in the FLAG-miP-FERMT3 pulldown compared to the EGFP (control) were analysed by STRING v.11.5 GO term enrichment analysis [65].

### Functional assays

#### Cell proliferation and cell area

Twenty-four hours post adenoviral transduction, cells were seeded at a density of 5,000 cells per well into a 96-well plate (day 0). After 3hr of attachment, cells were incubated and imaged for up to 5 days using Incucyte S3 live-cell imaging analysis system (Sartorius, Göttingen, Germany). Live cell count and average cell area were quantified by AI cell health analysis module (Segmentation sensitivity 0.5, Score threshold 0.2) within the Incucyte software.

#### Fluorescent Ubiquitination-based Cell Cycle Indicator (FUCCI) cell tracking

Cells expressing FUCCI were transduced with adenoviruses as described above. Similar to cell proliferation assay, cells were seeded and cultured at a density of 5000 cells per well into a 96-well plate (day 0). Images were acquired every hour using Incucyte S3 live-cell imaging and analysis system (Sartorius, Göttingen, Germany).

#### EdU incorporation

EdU assay was performed using Click-iT EdU Cell Proliferation Kit (Cat. # C10338; ThermoFisher Scientific; Darmstadt, Germany) according to the manufacturer’s instructions. Briefly, cells were incubated with 10 µmol/L EdU for 2 hours. Cells were then fixed with 4% PFA for 15 minutes and permeabilized with 0.5% Triton X-100 for 20 minutes at room temperature. Following two washes with 3% bovine serum albumin in PBS, cells were incubated with Click-iT® reaction cocktail for 30 minutes in the dark. The total number of cells and EdU positive cells were quantified using the Incucyte S3 live-cell imaging and analysis system (Sartorius, Göttingen, Germany). Results were shown as percent of EdU+/total number of cells.

### RNA isolation and next-generation RNA sequencing

Four days after adenoviral transduction, cells were lysed for 45-60 minutes at 4°C in a lysis buffer containing Tris-HCl pH7.5 (50mmol/L, Applichem GmbH; Darmstadt, Germany), NaCl (150mmol/L, Merck; Darmstadt, Germany), MgCl_2_ (10mmol/L, Invitrogen, Schwerte, Germany), NP-40 (1%, Merck; Darmstadt, Germany) dissolved in UltraPure DNase/RNase-free Distilled water (Invitrogen, Darmstadt, Germany) and supplemented with NaPPi (10mmol/L, Merck; Darmstadt, Germany), NaF (20mmol/L, Applichem GmbH; Darmstadt, Germany), okadaic acid (10nmol/L, LC laboratory, Massachusetts, USA), Na_3_VO_4_ (2mmol/L, Sigma-Aldrich; Darmstadt, Germany), PIM, (12µl/ml, Sigma-Aldrich; Darmstadt, Germany), Phenylmethylsulfonyl fluoride (4µl/ml, Carl Roth, Karlsruhe, Germany), SUPERase In RNase Inhibitor (200U/ml, Invitrogen, Darmstadt, Germany), TURBO DNaseI (25U/ml, Invitrogen, Darmstadt, Germany). The cell lysate was passed 7–10 times through a 1 mL syringe fitted with a 26G needle to ensure complete disruption of cellular membranes. The suspension was then centrifuged at 13,000 rpm for 10 minutes at 4 °C. The resulting clear supernatant was carefully transferred to a new 1.5 mL microcentrifuge tube for RNA purification. Total RNA was purified using the miRNeasy Micro kit (QIAGEN; Hilden, Germany), according to the manufacturer’s instructions.

RNA and library preparation integrity were verified with LabChip Gx Touch (Perkin Elmer). RNA amounts were normalized and 500ng/1µg of total RNA was used as input for SMARTer Stranded Total RNA Sample Prep Kit HI Mammalian (Takara Bio). Sequencing was performed on the NextSeq2000 platform (Illumina) using P3 flowcell with 72bp single-end setup. Trimmomatic version 0.39 was employed to trim reads after a quality drop below a mean of Q15 in a window of 5 nucleotides and keeping only filtered reads longer than 15 nucleotides [66]. Reads were aligned versus Ensembl human genome version hg38 (Ensembl release 109) with STAR 2.7.10a [67]. Alignments were filtered to remove: duplicates with Picard 3.0.0 (Picard: A set of tools (in Java) for working with next generation sequencing data in the BAM format), multi-mapping, ribosomal, or mitochondrial reads. Gene counts were established with featureCounts 2.0.4 by aggregating reads overlapping exons on the correct strand excluding those overlapping multiple genes [68]. The raw count matrix was normalized with DESeq2 version 1.36.0 [69]. Contrasts were created with DESeq2 based on the raw count matrix. Genes were classified as significantly differentially expressed at average count > 5, multiple testing adjusted p-value < 0.05, and -0.585 ≤ log2FC ≥ 0.585. The Ensemble annotation was enriched with UniProt data. Differentially expressed genes were analyzed by STRING v.11.5 GO term enrichment analysis [65].

FERMT3 mRNA expression analysis in human mesenteric artery endothelial cells from 20-versus 80-year-old individuals was obtained from the publicly available dataset GSE214476 [33]. FERMT3 mRNA expression analysis of isolated cardiac endothelial cells from 2-versus 20-month-old mice was analyzed as previously described [37].

### Flow cytometry

Cells were seeded at a density of 100,000 cells in 6-cm dishes and harvested after 24 hours (day 1) or 120 hours (day 5). Cells were fixed with 4% PFA for 15 minutes at room temperature, then incubated with 20 μmol/L Hoechst 33342 (B2261, Sigma-Aldrich; Darmstadt, Germany) for 45 minutes at 37 °C. After a single PBS wash, cells were resuspended in PBS and analyzed by flow cytometry. At least 50,000 events per sample were recorded for quantification. Fluorescence was analyzed using a BD LSR II/Fortessa flow cytometer (BD Biosciences; Heidelberg, Germany), and data were processed using FlowJo Vx software (TreeStar, UK). Daily instrument calibration was performed with Cytometer Setup and Tracking beads (BD Biosciences; Heidelberg, Germany).

### Senescence-associated β-galactosidase activity

Senescence-associated β-galactosidase activity was assessed using a Cellular Senescence assay kit (Cat. # KAA002, Merck; Darmstadt, Germany) according to the manufacturer’s instructions. In brief, cells were fixed with fixing solution (Part No. 2004755, 15 minutes at room temperature) 7 days post adenoviral transduction. After washing 3 times with PBS, cells were incubated with 150 µL of freshly prepared X-gal solution (Part No. 2004756, Part No. 2004754, Part No. 2004752) at 37 °C overnight. Cells were visualized in phase contrast mode using a Zeiss Axio Observer microscope (Zeiss; Oberkochen, Germany) and analyzed with the ZEN software (Zeiss; Oberkochen, Germany). At least 200 cells per cell batch were imaged and quantified. Results were expressed as senescence-associated β-galactosidase positive cells/total number of cells.

### Proteasome Activity assay

Proteasome activity was measured using the Proteasome 20S Activity Assay Kit (MAK172, Merck; Darmstadt, Germany) according to the manufacturer’s instructions. Briefly, five days post-transduction, cells were seeded in 96-well plates at a density of 40,000 cells per well and allowed to adhere for 3 hours. Cells were imaged using the the Incucyte S3 live-cell imaging and analysis system (Sartorius; Göttingen, Germany) to determine the number of adherent cells per well as described above, followed by incubation with Proteasome Assay Loading Solution overnight at 37 °C. Fluorescence intensity was measured at λ_ex_ = 485 nm and λ_em_ = 535 nm, and background fluorescence (medium without cells) was subtracted from each sample fluorescence value. Proteasome activity was expressed as corrected fluorescence intensity normalized to the total number of cells (proteasome activity per cell).

### Telomere length

Telomere length was quantified using a monochrome multiplexing qPCR method as described previously [70]. Briefly, genomic DNA was isolated from endothelial cells five days post-transduction using the DNeasy Blood and Tissue Kit (Cat. No. 69504, Qiagen, Hilden, Germany). For PCR, 20 ng genomic DNA per reaction was used. Primer pairs used for telomere (T) amplification were telomere forward primer: 5′-ACACTAAGGTTTGGGTTTGGGTTTGGGTTTGGGTTAGTGT-3′ and telomere reverse primer: 5′–TGTTAGGTATCCCTATCCCTATCCCTATCCCTATCCCTAACA–3′. As single copy gene reference (S), human β-globin was amplified at 88 °C using the primers β-globin forward primer: 5′–CGGCGGCGGGCGGCCGGGGCTGGGCGGCTTCATCCACGTTCACCTTG–3′ and β-globin reverse primer: 5′–GCCCGGCCCGCCGCGCCCGTCCCGCCGGAGGAGAAGTCTGCCGTT–3′. All reactions were carried out in quadruplicate. Relative telomere length was calculated as the telomere-to-single-copy gene (T/S) ratio using the Pfaffl method [71] and normalized to the mean values of control samples.

### Collection of conditioned media and monocyte adhesion assay

Forty-eight hours after adenoviral transduction, the culture media was replaced with ECGM2 and incubated with the cells for 96 hours. Conditioned media were subsequently used to treat naïve human endothelial cells for 24 hours. THP-1 monocytes were stained using 200 nmol/L CellTracker Red CMTPX dye (Cat. # C34552, ThermoFisher Scientific; Darmstadt, Germany) for 15 minutes at 37°C. After staining, cells were washed, counted, and resuspended in ECGM2 medium. Subsequently, 25,000 THP-1 cells were allowed to adhere to endothelial cells for 30 minutes. Non-adherent cells were removed by three thorough washes, and the number of adherent monocytes was quantified using the Incucyte S3 live-cell imaging and analysis system (Sartorius, Göttingen, Germany).

### Statistics

Results are presented as mean ± standard error of the mean. GraphPad Prism software (v. 10.1.2) was used to assess statistical significance. Differences between two groups were compared by two-tailed unpaired Student’s t-test. All experiments in which the effects of two variables were tested were analyzed by two-way ANOVA followed by the Šídák’s or Tukey’s multiple comparisons test. Differences were considered statistically significant when p-value < 0.05. Only exact significant p-values are reported.

## DECLARATIONS

### Ethics approval and consent to participate

The use of human material in this study complies with the principles outlined in the Declaration of Helsinki (World Medical Association, 2013), and the isolation of endothelial cells was approved in written form by the ethics committee of the Goethe-University.

### Consent for publication

Not applicable

### Availability of data and materials

Further information and requests for resources and reagents should be directed to and will be fulfilled by Dr. Mauro Siragusa. Data that support the findings of this study are available as part of the manuscript. Raw data can be accessed at the following repositories:

Mass spectrometry-based proteomics data: ProteomeXchange Consortium via the MassIVE partner repositories (http://proteomecentral.proteomexchange.org) with the following identifiers:

Microprotein miP-FERMT3: PXD069811 (MassIVE ID: MSV000099566)

Reviewer account details:

Bulk RNA-seq data are deposited in Gene Expression Omnibus (GEO) under the accession number GSE310464.

### Authors’ contributions

Manav Raheja, Beyza Güven, Witold Szymanski, Stefan Günther, Carsten Kuenne, Vladislav Rakultsev, Marta Segarra and Mauro Siragusa acquired, analyzed and interpreted the data. Manav Raheja drafted the manuscript. Johannes Graumann and Ingrid Fleming interpreted the results. Mauro Siragusa conceived the study, designed experiments and drafted the manuscript. All authors revised and approved the submitted version of the manuscript.

## Acknowledgements

The authors thank Sandrine Ngaha, Isabel Winter and Mechtild Piepenbrock-Gyamfi for expert technical assistance and Dr. Timo Frömel for support with flow cytometry analysis.

---

**Figure S1.**
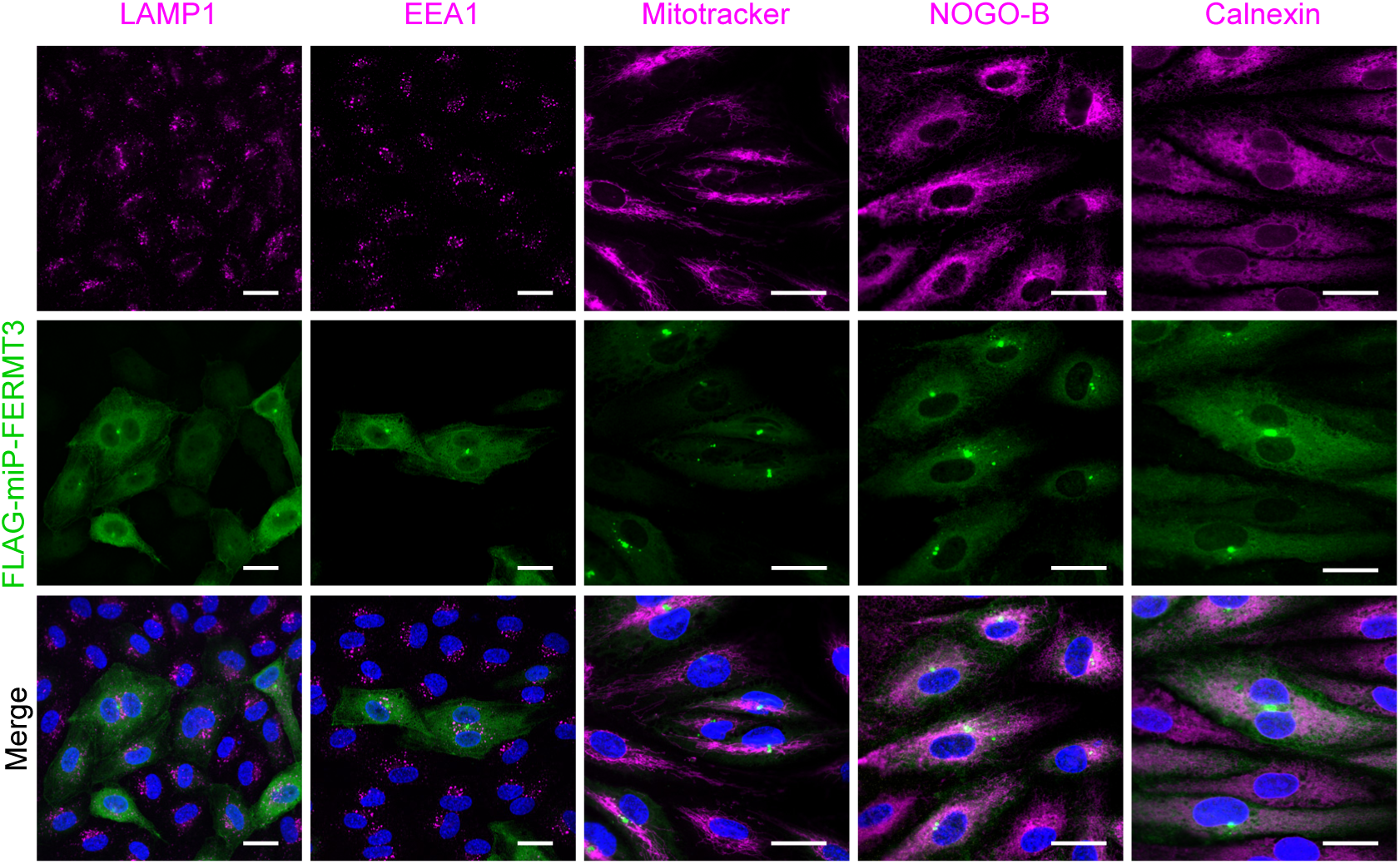
Subcellular localization of miP-FERMT3. Confocal images showing FLAG-miP-FERMT3 together with LAMP1 (lysosomes), EEA1 (early endosomes), Mitotracker (mitochondria), NOGO-B and calnexin (endoplasmic reticulum) in endothelial cells. Nuclei were stained with DAPI (blue). Similar results were obtained in 3-5 independent cell batches. Scale bar = 25 µm.

## REFERENCES

1. Slavoff SA, Heo J, Budnik BA, Hanakahi LA, Saghatelian A. A human short open reading frame (sORF)-encoded polypeptide that stimulates DNA end joining. J Biol Chem. 2014;289:10950–7. doi:10.1074/jbc.C113.533968.

2. Saghatelian A, Couso JP. Discovery and characterization of smORF-encoded bioactive polypeptides. Nat Chem Biol. 2015;11:909–16. doi:10.1038/nchembio.1964.

3. van Heesch S, Witte F, Schneider-Lunitz V, Schulz JF, Adami E, Faber AB, et al. The translational landscape of the human heart. Cell. 2019;178:242–260.e29. doi:10.1016/j.cell.2019.05.010.

4. Chothani S, Ruiz-Orera J, Tierney JAS, Clauwaert J, Deutsch EW, Alba MM, et al. An expanded reference catalog of translated open reading frames for biomedical research. bioRxiv 2025. doi:10.1101/2025.07.03.662928.

5. Mudge JM, Ruiz-Orera J, Prensner JR, Brunet MA, Calvet F, Jungreis I, et al. Standardized annotation of translated open reading frames. Nat Biotechnol. 2022;40:994–9. doi:10.1038/s41587-022-01369-0.

6. Siragusa M, Graumann J, Kuenne C, Günther S, Jeratsch S, Güven B, et al. High-throughput proteo-genomic identification of functional endothelial cell microproteins encoded by non-canonical small open reading frames. bioRxiv 2024. doi:10.21203/rs.3.rs-4194943/v1.

7. Chothani SP, Adami E, Widjaja AA, Langley SR, Viswanathan S, Pua CJ, et al. A high-resolution map of human RNA translation. Mol Cell. 2022;82:2885–2899.e8. doi:10.1016/j.molcel.2022.06.023.

8. Chen J, Brunner A-D, Cogan J, Nunez KJ, Fields AP, Adamson B, et al. Pervasive functional translation of noncanonical human open reading frames. Science 2020. doi:10.1126/science.aay026.

9. Anderson DM, Anderson KM, Chang C-L, Makarewich CA, Nelson BR, McAnally JR, et al. A micropeptide encoded by a putative long noncoding RNA regulates muscle performance. Cell. 2015;160:595–606. doi:10.1016/j.cell.2015.01.009.

10. Brito-Estrada O, Kuwabara Y, Gibson AM, Hassel KR, Kamradt ML, Verry JP, et al. DWORF gene therapy improves cardiac calcium handling and mitochondrial function. Circ Res. 2025;137:1072–88. doi:10.1161/CIRCRESAHA.125.326550.

11. M. Anderson D, A. Makarewich C, M. Anderson K, M. Shelton J, Bezprozvannaya S, Bassel-Duby R, N. Olson E. Widespread control of calcium signaling by a family of SERCA-inhibiting micropeptides. Sci Signal. 2016;9:1–6. doi:10.1126/scisignal.aaj1460.

12. Matsumoto A, Pasut A, Matsumoto M, Yamashita R, Fung J, Monteleone E, et al. mTORC1 and muscle regeneration are regulated by the LINC00961-encoded SPAR polypeptide. Nature. 2017;541:228–32. doi:10.1038/nature21034.

13. Li B, Zhang Z, Wan C. Identification of microproteins in Hep3B cells at different cell cycle stages. J Proteome Res. 2022;21:1052–60. doi:10.1021/acs.jproteome.1c00926.

14. Jayatissa A, Jaunbocus N, Erkalo B, Jiang K, Zheng S-J, Su H, et al. The ERVK3-1 microprotein interacts with the HUSH complex. Biochemistry. 2025;64:3372–81. doi:10.1021/acs.biochem.5c00023.

15. Wright BW, Yi Z, Weissman JS, Chen J. The dark proteome: translation from noncanonical open reading frames. Trends Cell Biol. 2022;32:243–58. doi:10.1016/j.tcb.2021.10.010.

16. D’Lima NG, Ma J, Winkler L, Chu Q, Loh KH, Corpuz EO, et al. A human microprotein that interacts with the mRNA decapping complex. Nat Chem Biol. 2017;13:174–80. doi:10.1038/nchembio.2249.

17. L. Hopper J, Begum N, Smith L, A. Hughes T. The role of PSMD9 in human disease: future clinical and therapeutic implications. AIMS Molecular Science. 2015;2:476–84. doi:10.3934/molsci.2015.4.476.

18. Hori T, Osaka Fumio, Chiba T, Miyamoto C, Okabayashi K, Shimbara N, et al. Covalent modification of all members of human cullin family proteins by NEDD8. Oncogene. 1999;18:6829–34. doi:10.1038/sj.onc.1203093.

19. Marzano F, Guerrini L, Pesole G, Sbisà E, Tullo A. Emerging roles of TRIM8 in health and disease. Cells 2021. doi:10.3390/cells10030561.

20. Caratozzolo MF, Micale L, Turturo MG, Cornacchia S, Fusco C, Marzano F, et al. TRIM8 modulates p53 activity to dictate cell cycle arrest. Cell Cycle. 2012;11:511–23. doi:10.4161/cc.11.3.19008.

21. Fischer M. Gene regulation by the tumor suppressor p53 - The omics era. Biochim Biophys Acta Rev Cancer. 2024;1879:189111. doi:10.1016/j.bbcan.2024.189111.

22. Hafner A, Bulyk ML, Jambhekar A, Lahav G. The multiple mechanisms that regulate p53 activity and cell fate. Nat Rev Mol Cell Biol. 2019;20:199–210. doi:10.1038/s41580-019-0110-x.

23. Keenan AB, Torre D, Lachmann A, Leong AK, Wojciechowicz ML, Utti V, et al. ChEA3: transcription factor enrichment analysis by orthogonal omics integration. Nucleic Acids Res. 2019;47:W212–W224. doi:10.1093/nar/gkz446.

24. Pandit B, Halasi M, Gartel AL. p53 negatively regulates expression of FoxM1. Cell Cycle. 2009;8:3425–7. doi:10.4161/cc.8.20.9628.

25. Barsotti AM, Prives C. Pro-proliferative FoxM1 is a target of p53-mediated repression. Oncogene. 2009;28:4295–305. doi:10.1038/onc.2009.282.

26. Piehl M, Tulu US, Wadsworth P, Cassimeris L. Centrosome maturation: measurement of microtubule nucleation throughout the cell cycle by using GFP-tagged EB1. Proc Natl Acad Sci U S A. 2004;101:1584–8. doi:10.1073/pnas.0308205100.

27. Lee M, Rhee K. Determination of mother centriole maturation in CPAP-depleted cells using the ninein Antibody. Endocrinol Metab (Seoul). 2015;30:53–7. doi:10.3803/EnM.2015.30.1.53.

28. Guarguaglini G, Duncan PI, Stierhof YD, Holmström T, Duensing S, Nigg EA. The forkhead-associated domain protein Cep170 interacts with Polo-like kinase 1 and serves as a marker for mature centrioles. Mol Biol Cell. 2005;16:1095–107. doi:10.1091/mbc.E04-10-0939.

29. Huang N, Xia Y, Zhang D, Wang S, Bao Y, He R, et al. Hierarchical assembly of centriole subdistal appendages via centrosome binding proteins CCDC120 and CCDC68. Nat Commun. 2017;8:15057. doi:10.1038/ncomms15057.

30. Pihan GA. Centrosome dysfunction contributes to chromosome instability, chromoanagenesis, and genome reprograming in cancer. Front Oncol. 2013;3:277. doi:10.3389/fonc.2013.00277.

31. Khurana A, Chadha Y, Schmoller KM. Too big not to fail: Different paths lead to senescence of enlarged cells. Mol Cell. 2023;83:3946–7. doi:10.1016/j.molcel.2023.10.024.

32. Tao W, Yu Z, Han J-DJ. Single-cell senescence identification reveals senescence heterogeneity, trajectory, and modulators. Cell Metab. 2024;36:1126–1143.e5. doi:10.1016/j.cmet.2024.03.009.

33. Hu J, Leisegang MS, Looso M, Drekolia M-K, Wittig J, Mettner J, et al. Disrupted binding of cystathionine γ-lyase to p53 promotes endothelial senescence. Circ Res. 2023;133:842–57. doi:10.1161/CIRCRESAHA.123.323084.

34. Han Z, Wang K, Ding S, Zhang M. Cross-talk of inflammation and cellular senescence: a new insight into the occurrence and progression of osteoarthritis. Bone Res. 2024;12:69. doi:10.1038/s41413-024-00375-z.

35. Freund A, Orjalo AV, Desprez P-Y, Campisi J. Inflammatory networks during cellular senescence: causes and consequences. Trends Mol Med. 2010;16:238–46. doi:10.1016/j.molmed.2010.03.003.

36. Ding Y, Zuo Y, Zhang B, Fan Y, Xu G, Cheng Z, et al. Comprehensive human proteome profiles across a 50-year lifespan reveal aging trajectories and signatures. Cell. 2025;188:5763–5784.e26. doi:10.1016/j.cell.2025.06.047.

37. Wagner UGJ, Tombor LS, Malacarne PF, Kettenhausen L-M, Panthel J, Kujundzic H, et al. Aging impairs the neurovascular interface in the heart. Science 2023. doi:10.1126/science.ade4961.

38. Takasugi M, Nonaka Y, Takemura K, Yoshida Y, Stein F, Schwarz JJ, et al. An atlas of the aging mouse proteome reveals the features of age-related post-transcriptional dysregulation. Nat Commun. 2024;15:8520. doi:10.1038/s41467-024-52845-x.

39. Makarewich CA. The hidden world of membrane microproteins. Exp Cell Res. 2020;388:111853. doi:10.1016/j.yexcr.2020.111853.

40. Rocha AL, Pai V, Perkins G, Chang T, Ma J, Souza EV de, et al. An inner mitochondrial membrane microprotein from the SLC35A4 upstream ORF Regulates Cellular Metabolism. J Mol Biol. 2024;436:168559. doi:10.1016/j.jmb.2024.168559.

41. Zhang S, Guo Y, Fidelito G, Robinson DRL, Liang C, Lim R, et al. LINC00116-encoded microprotein mitoregulin regulates fatty acid metabolism at the mitochondrial outer membrane. iScience. 2023;26:107558. doi:10.1016/j.isci.2023.107558.

42. Sangith N, Srinivasaraghavan K, Sahu I, Desai A, Medipally S, Somavarappu AK, et al. Discovery of novel interacting partners of PSMD9, a proteasomal chaperone: Role of an Atypical and versatile PDZ-domain motif interaction and identification of putative functional modules. FEBS Open Bio. 2014;4:571–83. doi:10.1016/j.fob.2014.05.005.

43. Faull SV, Lau AMC, Martens C, Ahdash Z, Hansen K, Yebenes H, et al. Structural basis of Cullin 2 RING E3 ligase regulation by the COP9 signalosome. Nat Commun. 2019;10:3814. doi:10.1038/s41467-019-11772-y.

44. Wei C-H, Weng C-W, Wu C-Y, Chen H-Y, Chang Y-H, Chang G-C, Chen JJW. E3 ligase TRIM8 suppresses lung cancer metastasis by targeting MYOF degradation through K48-linked polyubiquitination. Cell Death Dis. 2025;16:88. doi:10.1038/s41419-025-07421-6.

45. Sarikas A, Hartmann T, Pan Z-Q. The cullin protein family. Genome Biol 2011. doi:10.1186/gb-2011-12-4-220.

46. Caratozzolo MF, Marzano F, Abbrescia DI, Mastropasqua F, Petruzzella V, Calabrò V, et al. TRIM8 Blunts the Pro-proliferative Action of ΔNp63α in a p53 Wild-Type Background. Front Oncol. 2019;9:1154. doi:10.3389/fonc.2019.01154.

47. S Pellegata N, J Antoniono R, Redpath JL, J Stanbridge E. DNA damage and p53-mediated cell cycle arres_A reevaluation. Proc Natl Acad Sci U S A. 1996;93:15209–14. doi:10.1073/pnas.93.26.15209.

48. Macedo JC, Vaz S, Bakker B, Ribeiro R, Bakker PL, Escandell JM, et al. FoxM1 repression during human aging leads to mitotic decline and aneuploidy-driven full senescence. Nat Commun. 2018;9:2834. doi:10.1038/s41467-018-05258-6.

49. Wang I-C, Chen Y-J, Hughes DE, Ackerson T, Major ML, Kalinichenko VV, et al. FoxM1 regulates transcription of JNK1 to promote the G1/S transition and tumor cell invasiveness. J Biol Chem. 2008;283:20770–8. doi:10.1074/jbc.M709892200.

50. Smirnov A, Panatta E, Lena A, Castiglia D, Di Daniele N, Melino G, Candi E. FOXM1 regulates proliferation, senescence and oxidative stress in keratinocytes and cancer cells. aging. 2016;8:1384–97. doi:10.18632/aging.100988.

51. Wonsey RD, Wonsey DR, Follettie MT. Loss of the forkhead transcription factor FoxM1 causes centrosome amplification and mitotic catastrophe. Cancer Res. 2005;65:5181–9. doi:10.1158/0008-5472.CAN-04-4059.

52. Theile L, Li X, Dang H, Mersch D, Anders S, Schiebel E. Centrosome linker diversity and its function in centrosome clustering and mitotic spindle formation. EMBO J. 2023;42:e109738. doi:10.15252/embj.2021109738.

53. Fry AM, Meraldi P, Nigg EA. A centrosomal function for the human Nek2 protein kinase, a member of the NIMA family of cell cycle regulators. EMBO J. 1998;17:470–81. doi:10.1093/emboj/17.2.470.

54. Wu Q, Li B, Le Liu, Sun S, Sun S. Centrosome dysfunction: a link between senescence and tumor immunity. Signal Transduct Target Ther. 2020;5:107. doi:10.1038/s41392-020-00214-7.

55. Fleming I, Fisslthaler B, Dixit M, Busse R. Role of PECAM-1 in the shear-stress-induced activation of Akt and the endothelial nitric oxide synthase (eNOS) in endothelial cells. J Cell Sci. 2005;118:4103–11. doi:10.1242/jcs.02541.

56. Busse R, Lamontagne D. Endothelium-derived bradykinin is responsible for the increase in calcium produced by angiotensin-converting enzyme inhibitors in human endothelial cells. Naunyn-Schmiedeberg’s Arch Pharmacol. 1991;344:126–9. doi:10.1007/BF00167392.

57. Siragusa M, Fröhlich F, Park EJ, Schleicher M, Walther TC, Sessa WC. Stromal cell-derived factor 2 is critical for Hsp90-dependent eNOS activation. Sci Signal. 2015;8:ra81. doi:10.1126/scisignal.aaa2819.

58. Siragusa M, Thöle J, Bibli S-I, Luck B, Loot AE, Silva K de, et al. Nitric oxide maintains endothelial redox homeostasis through PKM2 inhibition. EMBO J. 2019;38:e100938. doi:10.15252/embj.2018100938.

59. Drekolia M-K, Mettner J, Wang D, Delgado Lagos F, Koch C, Hecker D, et al. Cystine import and oxidative catabolism fuel vascular growth and repair via nutrient-responsive histone acetylation. Cell Metab 2025. doi:10.1016/j.cmet.2025.10.003.

60. Llaó-Cid C, Peguera B, Kobialka P, Decker L, Vogenstahl J, Alivodej N, et al. Vascular FLRT2 regulates venous-mediated angiogenic expansion and CNS barriergenesis. Nat Commun. 2024;15:10372. doi:10.1038/s41467-024-54570-x.

61. Chen F, Tillberg PW, Boyden ES. Optical imaging. Expansion microscopy. Science. 2015;347:543–8. doi:10.1126/science.1260088.

62. Hughes CS, Moggridge S, Müller T, Sorensen PH, Morin GB, Krijgsveld J. Single-pot, solid-phase-enhanced sample preparation for proteomics experiments. Nat Protoc. 2019;14:68–85. doi:10.1038/s41596-018-0082-x.

63. Demichev V, Messner CB, Vernardis SI, Lilley KS, Ralser M. DIA-NN: neural networks and interference correction enable deep proteome coverage in high throughput. Nat Methods. 2020;17:41–4. doi:10.1038/s41592-019-0638-x.

64. Ritchie ME, Phipson B, Di Wu, Hu Y, Law CW, Shi W, Smyth GK. limma powers differential expression analyses for RNA-sequencing and microarray studies. Nucleic Acids Res. 2015;43:e47. doi:10.1093/nar/gkv007.

65. Szklarczyk D, Gable AL, Nastou KC, Lyon D, Kirsch R, Pyysalo S, et al. The STRING database in 2021: customizable protein-protein networks, and functional characterization of user-uploaded gene/measurement sets. Nucleic Acids Res. 2021;49:D605–D612. doi:10.1093/nar/gkaa1074.

66. Bolger AM, Lohse M, Usadel B. Trimmomatic: a flexible trimmer for Illumina sequence data. Bioinformatics. 2014;30:2114–20. doi:10.1093/bioinformatics/btu170.

67. Dobin A, Davis CA, Schlesinger F, Drenkow J, Zaleski C, Jha S, et al. STAR: ultrafast universal RNA-seq aligner. Bioinformatics. 2013;29:15–21. doi:10.1093/bioinformatics/bts635.

68. Liao Y, Smyth GK, Shi W. featureCounts: an efficient general purpose program for assigning sequence reads to genomic features. Bioinformatics. 2014;30:923–30. doi:10.1093/bioinformatics/btt656.

69. Love MI, Huber W, Anders S. Moderated estimation of fold change and dispersion for RNA-seq data with DESeq2. Genome Biol. 2014;15:550. doi:10.1186/s13059-014-0550-8.

70. Cawthon RM. Telomere length measurement by a novel monochrome multiplex quantitative PCR method. Nucleic Acids Res. 2009;37:e21. doi:10.1093/nar/gkn1027.

71. Pfaffl MW. A new mathematical model for relative quantification in real-time RT-PCR. Nucleic Acids Res. 2001;29:e45. doi:10.1093/nar/29.9.e45.

